# Manipulating the murine *Lgr5* locus using a rapid, efficient and flexible CRISPR/Cas9 pipeline

**DOI:** 10.1101/2020.12.31.424946

**Authors:** Jan Reichmuth, Johannes vom Berg, Michael Brügger, George Hausmann, Tomas Valenta, Konrad Basler

## Abstract

The maintenance of adult epithelial tissues, such as the lining of the intestine, depends on their periodical renewal. This is achieved through small populations of adult stem cells. Lgr5 serves as a marker for these cells. Here we report a novel non-variegated Lgr5 mouse model, which was generated via the CRISPR/Cas9 system. We show that this *Lgr5-2A-CreERT2-2A-mOrange2* mouse line can be used for lineage tracing, as well as for directing gene expression to Lgr5^+^ cells. The introduction of the transgene affects neither the expression, nor the function of endogenous *Lgr5*. Therefore, this new tool will serve to mark and manipulate intestinal stem cells to gain new biological insights.

## Introduction

The transmembrane receptor Lgr5 (Leucin-rich repeat-containing G protein-coupled receptor 5) was identified as a marker of adult stem cells in many epithelial tissues. Lgr5^+^ stem cells were first found in the intestinal epithelium, a fast renewing tissue with a turnover rate of roughly 5-7 days (Barker et al., 2007). Since then, Lgr5^+^ stem cells have been identified in various other tissues like hair follicle, stomach, mammary glands and ovaries (Jaks et al., 2008; Barker et al., 2010; Plaks et al., 2013; Ng et al., 2014). Additionally, it was shown in liver, pancreas and stomach, that quiescent Lgr5^+^ stem cells can get activated and start proliferating upon injury to support tissue regeneration (Huch et al., 2013a, 2013b; Leushacke et al., 2017). Additionally, colorectal and gastric cancer, intestinal adenomas or squamous cell carcinomas have been reported to originate from Lgr5^+^ stem cells upon acquisition of oncogenic mutations (Barker et al., 2009; Schepers et al., 2012; Li et al., 2016; Melo et al., 2017; Leushacke et al., 2017; Huang et al., 2017). Accordingly, Lgr5 expression is often significantly upregulated in various subtypes of cancers and Lgr5^+^ cancer stem cells contribute to tumour growth (Junttila et al., 2015; Gong et al., 2016; Cortina et al., 2017; Shimokawa et al., 2017).

The Wnt/β-catenin signalling pathway is essential to drive epithelial proliferation and thus maintain tissue homeostasis. *Lgr5* is a Wnt target gene and the Lgr5 protein was shown to potentiate Wnt-signalling (de Lau et al., 2011). The ubiquitin ligases Znrf3 and Rnf43 mark the Frizzled receptors for degradation (Hao et al., 2012). Binding of R-spondin to Lgr5 and Znrf3/Rnf43 inhibits this process and thus boosts the output of canonical Wnt-signalling. Lgr5-based mouse models therefore present an excellent tool to study homeostasis, regeneration and disease, especially in epithelial tissues.

Several Lgr5 based mouse models have been described, with the Lgr5-CreERT2-IRES-EGFP (further referred to as Lgr5^UT^) reporter line currently being the most frequently used one (Barker et al., 2007; Tian et al., 2011; Leushacke et al., 2017). However, this model is homozygous lethal and oftentimes displays variegated expression levels.

Here we used the CRISPR/Cas9 system to manipulate the *Lgr5* locus without interfering with its endogenous protein function. With this approach we were able to generate a novel Lgr5 mouse model, which can be utilized to gain further insights into the role and molecular mechanisms of Lgr5^+^ stem cells. Furthermore, the provided homology repair template enables fast and flexible generation of other *Lgr5*-dependent mouse models.

## Results

To circumvent variegated expression of the transgene and causing a *Lgr5* null allele, we chose to manipulate the 3’ end of *Lgr5*. We decided to use a CRISPR-based approach, since the generation of mouse models using this method is faster than homologous recombination in mouse embryonic stem cells (Hsu et al., 2014). gRNAs targeting a site directly prior to the STOP codon of *Lgr5* were selected and tested in vitro to determine their cleavage potential (Supplementary Figure 1a).

To modify the *Lgr5* locus, Cas9 bound gRNA was injected into the male pronucleus of zygotes together with a homology-directed repair (HDR) DNA template (Figure 1a). The HDR template was designed with the Lgr5 homology arms, an in-frame 2A sequence and a multiple cloning site (MCS). The MCS allows maximum flexibility as different genes (or inserts) of interest can easily be cloned into it (Supplementary Figure 2). Accordingly, we generated a *CreERT2-2A-mOrange2* cassette, which was subsequently integrated at the 3’ end of *Lgr5* (Figure 1b). As a result, Lgr5-expressing cells are directly marked with the fluorophore mOrange2 and the inducible CreERT2 allows temporally controlled manipulation of LoxP based conditional alleles (e.g. lineage tracing or conditional knock-outs). 2A peptides, which cause ribosomal skipping during translation, separate Lgr5, the CreERT2 recombinase and mOrange2 (Kim et al., 2011). This prevents functional interference between the Lgr5 receptor and the introduced downstream coding sequences. Moreover, this setup ensures that the coding sequences introduced downstream of *Lgr5* are expressed at equal levels.

**Figure 1:**
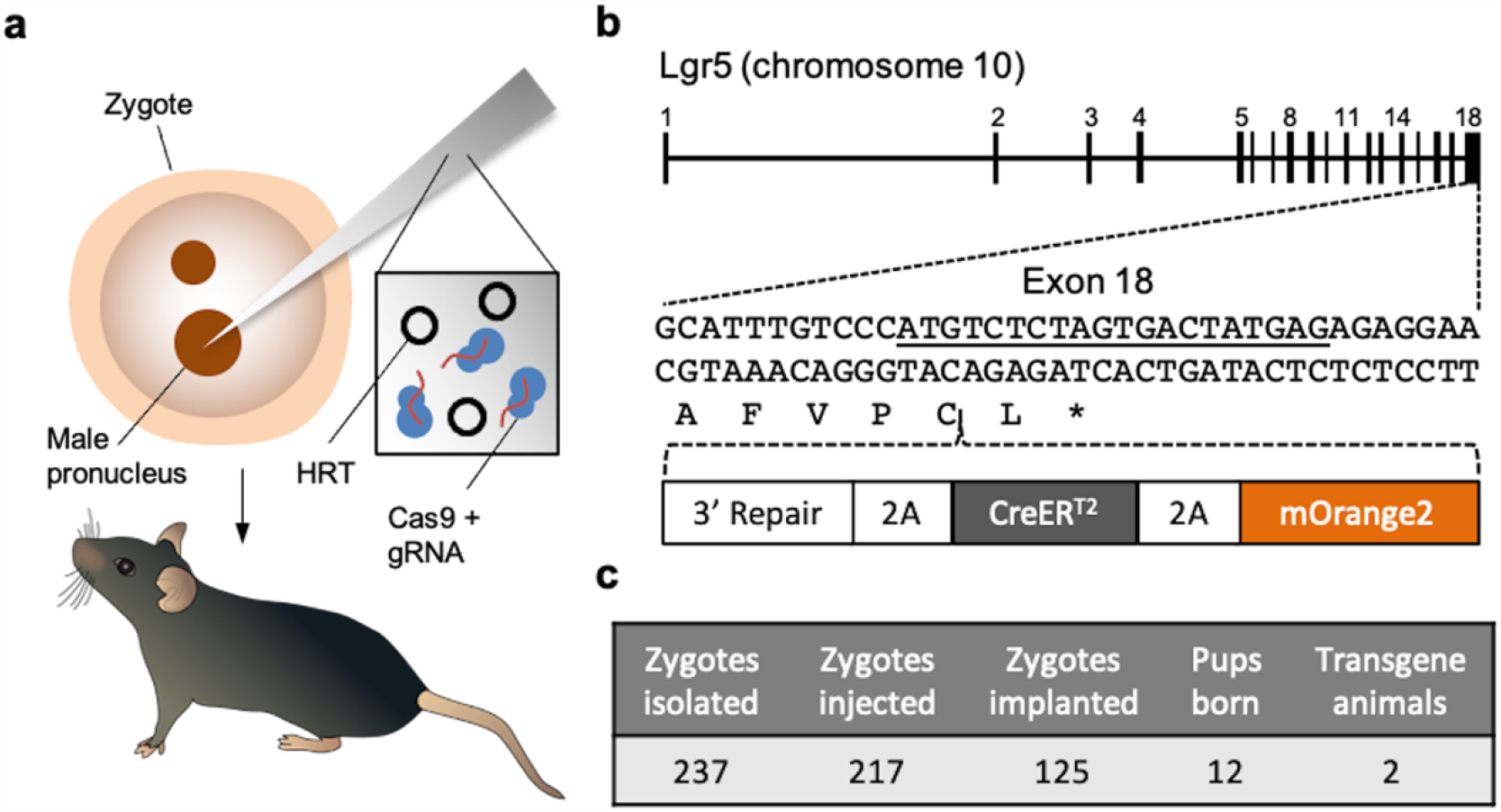
Schematic overview of the construct and integration efficacy. a) Schematic representation of the *Lgr5* endogenous locus with translational STOP depicted in detail. The DNA is targeted by CRISPR/Cas9 (gRNA annealing indicated by the underscore) prior to the STOP codon within the Cysteine (C). The introduced sequence contains a repair for the 3’ end, the Cre recombinase CreERT2 and the fluorophore mOrange2. The genes are separated by self-cleaving 2A sequences. b) Pronuclear microinjection: The homology directed repair (HDR) template is injected together with gRNA loaded Cas9 protein into the pronucleus fertilized murine zygote. The surviving zygotes are then transplanted into pseudopregnant females to carry out the pups. c) Injection statistics Lgr5^ZH^: Indicated are the isolated, injected and transplanted zygotes, as well as the number of born pups and if transgenes with correct integration were obtained.

In one round of injections 217 zygotes were injected and 125 of those zygotes were then transplanted into five pseudopregnant foster females (Figure 1c). Twelve pups were born and genotyping revealed two of those animals had heterozygous integrations of the *2A-CreERT2-2A-mOrange2* construct (Figure 1c, Supplementary Figure 1b). Heterozygous, as well as homozygous, *Lgr5-2A-CreERT2-2A-mOrange2* mice (henceforth referred to as Lgr5^ZH^) were healthy and fertile. A comparison between the colon and the duodenum of homozygous and heterozygous Lgr5^ZH^ animals revealed no morphological differences.

As a next step we functionally validated the Lgr5^ZH^ reporter model. Lgr5 expression is considered a marker of intestinal epithelial stem cells, which are located at the bottom of the crypt (Barker et al., 2007). Hence in Lgr5^ZH^ mice mOrange2 expression should also be detectable at the crypt bottom. As expected, in the small (Figure 2a) and large intestine (Figure 2c), the expression of mOrange2 was restricted to the bottom of the crypts, confirming the allele’s specificity for labelling only the stem cells. The homozygous Lgr5^ZH^ alleles expressed higher mOrange2 levels, as confirmed by fluorescence-activated cell sorting (FACS) (Figure 2b and b’ for small intestine, d and d’ for colon). Together the results show that mOrange2 is strongly expressed, effectively marking the Lgr5 stem cell population.

**Figure 2:**
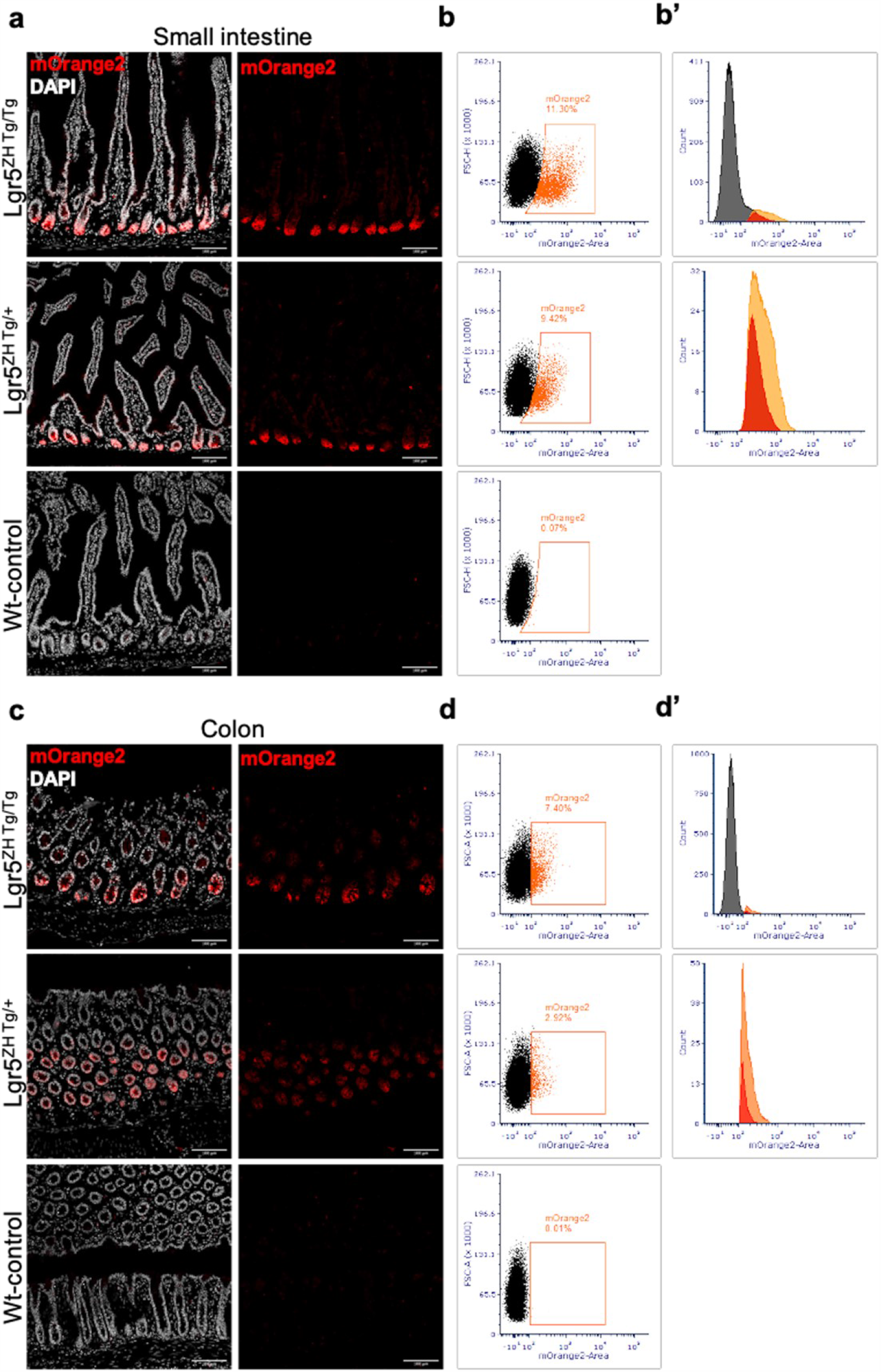
Functional validation of the mOrange2 fluorophore in Lgr5^ZH^ mice. mOrange2 staining of cryosections from OCT embedded small intestine a) or colon c) revealed restriction of mOrange2 expression at the bottom of the crypt in homozygous and heterozygous Lgr5^ZH^ animals. FACS analysis of isolated crypt cells found more mOrange2^+^ (percentage indicated in gating) cells in the small intestine b), and in the colon d). In both tissues the homozygous allele showed more mOrange2^+^ cells, as well as a higher signalling intensity within the mOrange^+^ population as the respective heterozygous allele. Overlay of the FACS analysis histograms for mOrange2 (b’, d’) indicates the difference between homozygous (orange) and heterozygous (red) Lgr5^ZH^ allele.

To validate CreERT2 efficacy, Lgr5^ZH^ mice were crossed to mice carrying the Cre-inducible *Rosa26-LSL-tdTomato* reporter allele (Madisen et al., 2010, Figure 3a). The *Lgr5*^*ZH*^; *Rosa26-LSL-tdTomato* animals then received a single tamoxifen injection to induce CreERT2-mediated recombination and multiple time points were evaluated. We examined recombination in the small intestine (Figure 3b) and the colon (Figure 3c). To quantify the recombination efficacy, crypts of at least five independent stretches of proximal, mid and distal intestinal epithelia were analysed per animal (n ≥ 2). Crypts were considered recombined when at least one cell within the crypt was expressing tdTomato. One day after induction, individual cells started to express tdTomato at the bottom of the crypts where the Lgr5+ stem cells are located. Quantification yielded tdTomato positive cells in more than 90 % of the crypts (147 of 159 analysed crypts) in the small intestine (Supplementary Figure 3a). At day five large tdTomato positive ribbons spanned the crypt and along the villi in the small intestine and 95 % of all crypts in the small intestine (397 of 416 analysed crypts) showed tdTomato expressing cells (Figure 3b, Supplementary Figure 3a). Thirty days after the induction, tdTomato expression was observed in 247 of 252 analysed crypts (98 %) and tdTomato positive ribbons appeared throughout the entire length of the villi in the small intestine (Figure 3b, Supplementary Figure 3a).

**Figure 3:**
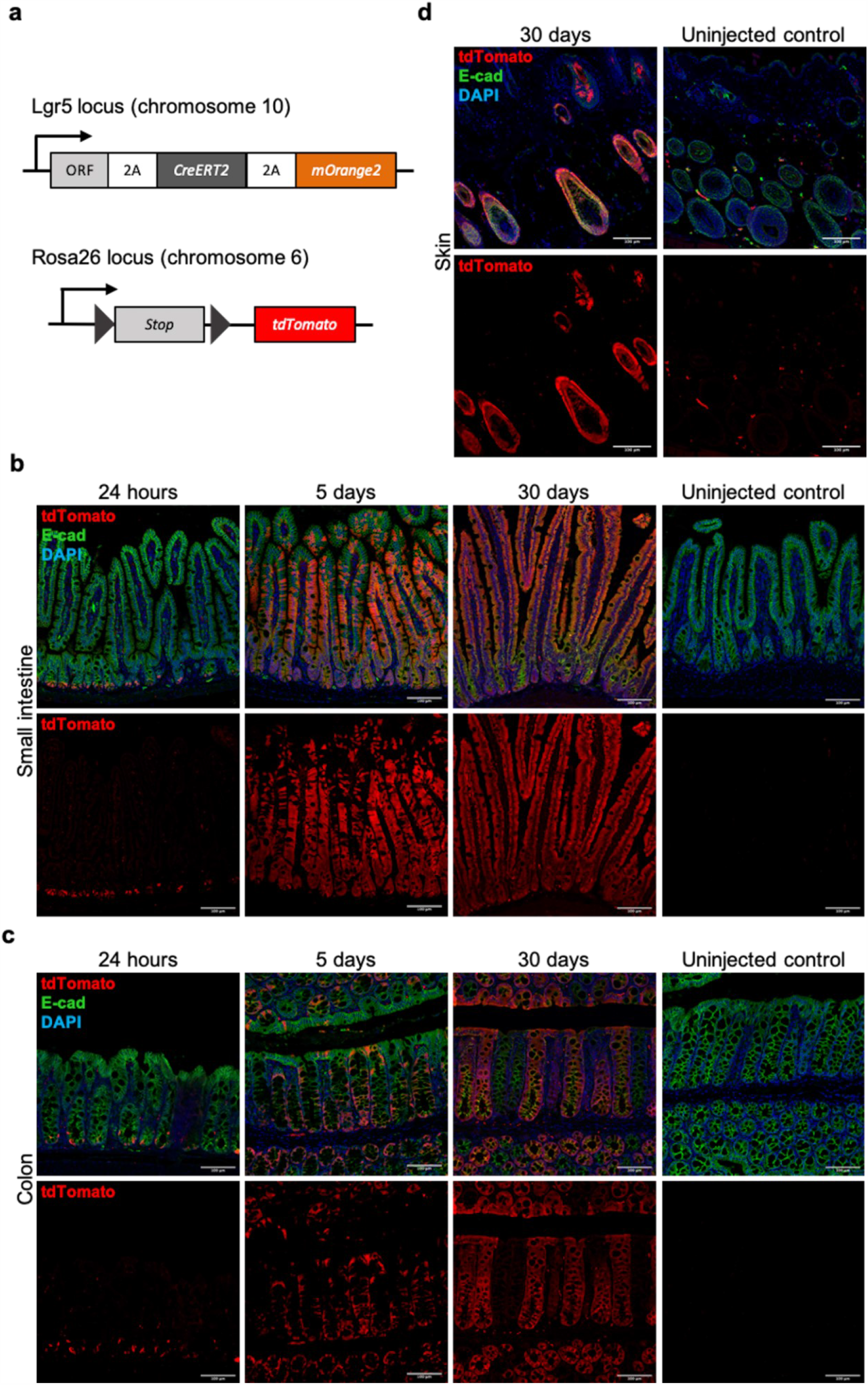
Functional assessment of the CreERT2 recombinase. Lgr5^ZH^ animals were combined with Rosa26-LSL-tdTomato animals to validate CreERT2 function. Induction by single tamoxifen injection causes the recombination determined by the expression of tdTomato, which was evaluated at different time points. In the small intestine b) one day after induction tdTomato^+^ cells started to appear at the bottom of the crypt. 5 days post injection large ribbons of tdTomato^+^ cells were noticed, covering almost the entire length of the villi. 30 days after induction whole small intestine was covered with tdTomato^+^ cells, leaving no unrecombined cells. In the colon c) tdTomato expression was found also one day after induction. 5 days after, the vast majority of bottom crypt cells were expressing tdTomato and most crypts showed emerging tdTomato^+^ ribbons. 30 days after induction large ribbons of tdTomato^+^ cells, covering entire crypts were observed. However, individual crypts were found to lack tdTomato. d) Recombination was also confirmed in anagen hair follicles cells in the skin.

In the colon, tdTomato expression was also found widespread throughout the crypts (Figure 3c). At day one after induction 67% of the crypts (86 of 127) in the colon showed tdTomato positive cells (Supplementary Figure 3b). At day five, there were scattered crypts which showed no tdTomato positive cells but 654 of 810 analysed colonic crypts (80 %) revealed recombined cells (Figure 3c, Supplementary Figure 3b). Thirty days after the induction recombination was found in 78 % of the colonic crypts (247 of 252 analysed crypts). Those crypts were entirely positive for tdTomato (Figure 3c, Supplementary Figure 3b).

It should be mentioned that in rare occasions also the uninjected *Lgr5*^*ZH*^; *Rosa26-LSL-tdTomato* control animals showed tdTomato positive patches in the small intestine and the colon (Supplementary Figure 3c and d). Those recombined patches were not found in *Rosa26-LSL-tdTomato* control animals without the *Lgr5*^*ZH*^-driver.

The functionality of CreERT2 was also tested in the hair follicles where Lgr5 positive cells proliferate during the anagen phase (Jaks et al., 2008). Thirty days after recombination of the *Lgr5*^*ZH*^; *Rosa26-LSL-tdTomato* animals tdTomato expression could be observed in the hair follicles (Figure 3d). This is in line with previously observed *Lgr5* expression patterns for anagen hair follicle proliferation (Jaks et al., 2008). The results confirm the functionality of the CreERT2 protein and show that a single injection of Tamoxifen is sufficient to achieve recombination in most of the Lgr5 positive stem cells in both intestine and hair follicles.

Next, we compared our *Lgr5*^*ZH*^ allele to the frequently used *Lgr5*^*UT*^ allele. Both alleles were combined with the *Rosa26-LSL-tdTomato* allele and recombination efficiency was assessed five days post injection in the intestine. For comparison only heterozygous *Lgr5*^*ZH*^ animals were used animals since the *Lgr5*^*UT*^ allele is homozygous not viable. The animals were examined five days after a single tamoxifen injection. With the *Lgr5*^*UT*^ allele 65 of the 175 analysed crypts (37 %) in the small intestine displayed tdTomatopositive cells, whereas in the *Lgr5*^*ZH*^ allele 95% of the crypts (397 of 416) showed tdTomato positive cells (Figure 4a and b). In the colon the recombination rate in general was lower than in the small intestine. However, with the *Lgr5*^*ZH*^ allele we could still observe tdTomato positive cells, whereas for the *Lgr5*^*UT*^ allele the majority (about 70%) of crypts remained unrecombined and did not show any tdTomato expression. To get a better overview and assess more crypts in the colon, transversal cuts through the crypts were performed (Figure 4e, f). Those sections confirmed the previous observations. The *Lgr5*^*UT*^ mice had recombined cells in 81 of the 304 analysed colonic crypts (about 30 %) compared to 654 of 810 crypts in the Lgr5^ZH^ mice (80 %) (Figure 4d). The results show that the *Lgr5*^*ZH*^ allele is an efficient Cre-driver.

**Figure 4:**
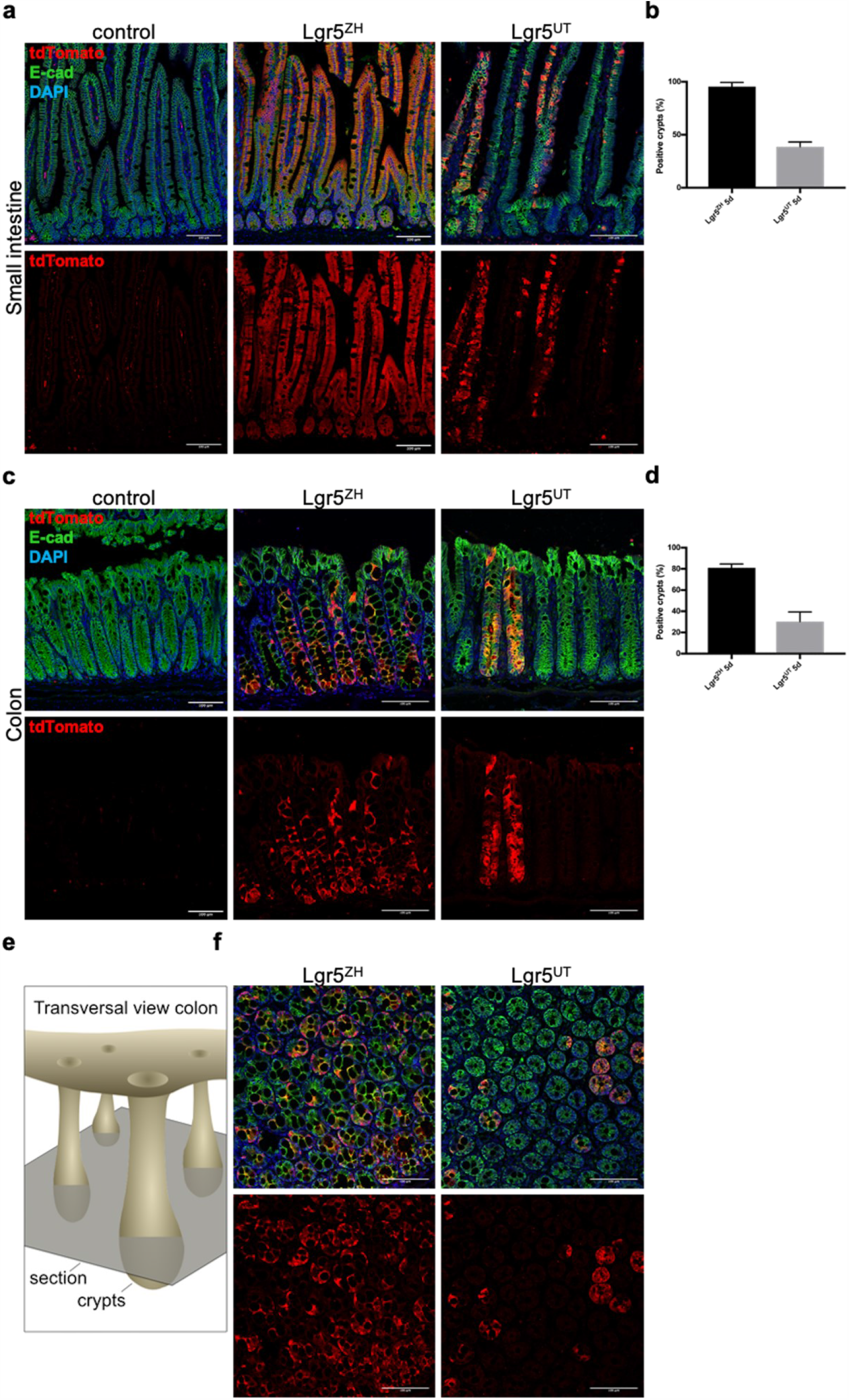
Side by side comparison of Lgr5^ZH^ and Lgr5^UT^. Lgr5^ZH^ and Lgr5 ^UT^ animals were combined with Rosa26-LSL-tdTomato animals. a) Five days after the induction Lgr5 ^UT^ showed large ribbons spanning the small intestine but not all crypts and villi were tdTomato^+^, whereas Lgr5^ZH^ was found to express tdTomato in almost all crypts. b) Quantification tdTomato^+^ crypts in the small intestine. Cryps were considered positive when at least one cell expressed tdTomato within the crypt. Minimum 4 independent stretches of crypts from proximal, mid and distal small intestinal epithelium were analyzed per animal (n ≥ 2). c) The colon mimicked the observed phenotype in the small intestine. TdTomato^+^ Ribbons were sparsely developed in Lgr5^UT^ animals and abundant in Lgr5^ZH^ animals. d) Quantification colon as indicated before. e) Schema transversal section. f) transversal sections of Lgr5^ZH^ and Lgr5 ^UT^ showed more recombination in Lgr5^ZH^.

## Discussion

The importance and universal use of Lgr5 as a marker for epithelial stem cells have become clear for more than one decade since its discovery (Barker et al., 2007). Here we describe a novel homozygous viable, non-variegated murine *Lgr5* reporter line. The strain combines a fluorophore to locate and mark the Lgr5^+^ stem cell population and CreERT2, it thus can be used for lineage tracing experiments as well as Lgr5-depended gene ablation/expression. We show that the fluorophore – mOrange2 – marks stem cells in the colon and small intestine. We also show that the CreERT2 is functional. Compared to the *Lgr5*^*UT*^ allele, we find that our CreERT2 line is more efficient, especially in the colon. With a single tamoxifen injection, we achieved close to 100% recombination of the LSL-tdTomato allele in the small intestine and up to 80% in colon. A likely explanation for the different efficiencies is the use of a 2A sequence rather than an IRES (Internal Ribosomal Entry Site) sequence. Contrary to an IRES where expression levels of secondary proteins are often lower, the 2A peptide guarantees that the expression levels of the downstream peptides are the same as those of Lgr5 (de Felipe et al., 2006). Utilizing an IRES sequence frequently leads to significantly lower expression of the downstream sequence (Mizuguchi et al., 2000).

An advantage of the 3’ integration and the use of the 2A peptide is the fact that both *Lgr5* alleles remain functional and we do not interfere with Lgr5 function. The *Lgr5*^*UT*^ allele is only heterozygous viable, whereas our allele is homozygous viable (Barker et al., 2007). In addition to the potential to boost recombination efficiency it also simplifies the genetics of crosses.

The Barker group more recently published a *Lgr5-2A-CreERT* mouse allele, which also used a 3’ integration at the endogenous *Lgr5* locus (Leushacke et al., 2017). We did not have access to this line for a comparison. However, the lines are likely to have similar CreERT2 expression and activity. As shown in Leushacke et al. (2017), such drivers can give new insights. A novel feature of our line is that we have also incorporated a direct marker, which will facilitate the identification and tracing of Lgr5 expressing cells. We also provide a detailed map of the HDR template, which can be used for the CRISPR-based generation of other novel *Lgr5* alleles (Supplement Figure 2).

Acknowledgements

We are grateful to all members of the Basler lab, in particular to J. Schopp and N. Doumpas, for valuable comments. For technical help we thank E. Escher and the Cytometry Facility of the University of Zurich. This work was supported by the Swiss National Science Foundation, the Swiss Cancer League, the University of Zurich Research Priority Program (URPP) “Translational Cancer Research” and the Kanton of Zürich. MB is supported by the Forschungskredit of the University of Zürich grant FK-19-074. TV is supported by Czech Science Foundation and is a fellow of the URPP Translational Cancer Research.

## Material and Methods

### Generation of mice

The *Lgr5-2A-CreERT2-mOrange2* mice were generated via pronuclear microinjection of Cas9 protein loaded gRNA and the plasmid harbouring the homology directed repair (HDR) template in fertilized zygotes.

The guide RNA (5’ CTCATAGTCACTAGAGACAT 3’) was selected using the gRNA prediction tool CRISPOR (http://crispor.tefor.net/) and tested *in vitro* (Sup Fig.1). For the *in vitro* test 400 ng column cleaned up PCR product was used per condition. Cas9NLS was loaded on the gRNA and incubated for 15 min to simulate a pronuclear injection mix as described below. Purified PRC DNA was digested for 1 h before run on gel electrophoresis.

To generate HDR donor plasmids, Lgr5 homology arms with 2A sequence and multiple cloning site were synthesized (Invitrogen). For the generation of the *Lgr5-2A-CreERT2-2A-mOrange2* cassette, CreERT2 was amplified from genomic DNA of VillinCreERT2 animals and combined with the 2A (annealed primers, Microsynth AG, Switzerland) and mOrange2 (cDNA). This was then transferred into the plasmid containing the homology arms.

For the preparation of RNP/HDR donor injection mixes, lyophilized crRNA and tracrRNA (Alt-R crispr, iDT) were resuspended in 1x microinjection buffer (TrisHCl 10 mM, pH 7.5, EDTA 0.1 mM) to a final concentration of 10µM. 1.84µl of tracr and crRNA were mixed with 10x injection buffer (5µl) and 0,5µl of streptococcus pyogenes Cas9 protein (Engen Cas9 NLS, 20µM, New England Biosciences) and subsequently incubated for 15 minutes at 37°C. After incubation, 500ng of HDR containing plasmid were added and the mix was diluted with ddH2O to a final volume of 50µl. To sediment any debris before injection, the final mix was spun at 21000g for 3 min at room temperature. Injection mix was kept at RT during the microinjection procedure.

Microinjection was performed at the transgenesis core of the University of Zurich, Institute of Laboratory Animal Science under cantonal veterinary office license nr 177-G. C57BL/6 mice at 3-4 weeks of age (Charles River Germany) were superovulated by intraperitoneal injection of 5 IU pregnant mare serum gonadotropin (Folligon, MSD Animal Health GmbH, Luzern, Switzerland) followed 48 h later by injection of 5 IU human chorionic gonadotropin (Pregnyl MSD Animal Health GmbH, Luzern, Switzerland). Mouse zygotes were obtained by mating C57BL/6 stud males with superovulated C57BL/6 females. Zygote microinjections, embryo culture and retransfer into pseudopregnant foster animals were performed according to standard transgenesis protocols (Harms et al., 2014). The obtained pups were biopsied and genotyped as described below. Founders were backcrossed to C57BL/6J for two generations.

### DNA preparation and genotyping

Tissue biopsies were lysed overnight at 55 °C in 500 µl buffer (100 mM Tris/HCL pH 7.5, 5 mM EDTA, 0.2% SDS, 200 mM NaCl, 10 ng Proteinase K) and DNA was isolated by Isopropanol precipitation and resuspended in 100 µl TE buffer. PCR was performed according to the GoTaq DNA Polymerase protocol (https://ch.promega.com/resources/protocols/product-information-sheets/g/gotaq-dna-polymerase-m300-protocol/), using 3 µl DNA as template. For primer information see Supplementary Figure 1c.

### Breeding

*Lgr5*^*ZH*^ and *Lgr5*^*UT*^ mice were bred to *Rosa26-LSL-tdTomato* (Ai14) mice (Madisen et al., 2010). The *Rosa26-LSL-tdTomato* mice were obtained from Jackson Labs (Ai14; stock no. 007914). The *Lgr5*^*UT*^ mice have been described previously (Barker et al., 2007). All mouse experiments were approved by the Veterinarian Office of Kanton Zürich (Switzerland) and were performed according to Swiss guidelines. Mice of >8 weeks were injected intraperitoneally with tamoxifen (80 mg/kg, Sigma) in sunflower oil to induce CreERT2-mediated recombination.

### Immunohistochemistry

The animals were euthanized and the duodenum and colon were dissected immediately after. The tissue was flushed with cold PBS, cut into pieces and then fixed in 4 % paraformaldehyde (PFA). For cryosections 30 min PFA fixation at room temperature (RT) was used before equilibrating it in 30 % sucrose in PBS over night at 4°C. The tissue was then incubated at 4°C for additional 2-4 h in a mixture of OCT (Tissue-Tek, #4583) and 30 % sucrose. Afterwards the tissue pieces were embedded in 100 % OCT and frozen at −80 °C, where they were stored until cutting (8 µm).

For paraffin sectioning, the tissue was fixed overnight at 4 °C in 4 % PFA and afterwards stepwise dehydrated in 30, 50 and 70 % EtOH in PBS (1 h each step on a rotator at RT) before embedding it in paraffin. 5 µm sections were cut and after deparaffination antigen retrieval was done by boiling the sections for 20 min in 1.6 mM citric acid and 8.4 mM trisodium citrate. The following primary antibodies were used for immunostaining: Rabbit anti-RFP (600-401-379, Rockland, 1:200), mouse anti E-cadherin (610181, BD Transduction Laboratories, 1:200). As secondary antibodies Alexa Fluor 555 (goat anti-rabbit, A32732, Life Technologies) and 647 (goat anti-mouse, A32728, Life Technologies) were used at a 1:400 dilution.

### Microscopy

For image acquisition a Zeiss LSM 710 and Leica SP8 inverse CLSM with ZEN software were used. Images were processed in Fiji (ImageJ). For the quantification of tdTomato^+^ crypts different sections from proximal to distal small intestine and colon were analysed. Crypts with at least one tdTomato^+^ cell were considered positive and counted as such. Ratios of tdTomato^+^ crypts to total crypt numbers were calculated.

### Crypt isolation, cell dissociation and FACS analysis

Colon and duodenum were dissected and flushed with PBS. Afterwards the tissues were opened longitudinally and washed again before cutting them in 1-2 mm pieces. The colon and duodenum pieces were washed by brief vortexing 5 times with 10 ml PBS, followed by 10-15 washes by pipetting up and down with 10 ml PBS until the supernatant is clear. Afterwards the crypts were liberated from the tissue by incubation (20 min duodenum, 40 min colon) on RT on a rotator in Gentle cell dissociation reagent (Stem cell technologies, #07174). Afterwards the dissociation reagent was removed and the tissue was resuspended in 10 ml 0.1% BSA in PBS by pipetting up and down. This was repeated 4 times (first fraction is discarded) and isolated crypts were passed through a 70 µm cell strainer (Falcon, 352350). The 3 remaining fractions were pooled and pelleted at 290 g for 4 min. The supernatant was discarded and the cells were dissociated for 5 min on 37°C using TrypLE Express (Thermo Fisher, #12604013). Immediately afterwards the TrypLE was inactivated, cells were pelleted and after resuspending in 1 ml PBS passed through a 35 µm cell strainer (Falcon, 352235). Single cells were then incubated on ice for 15 min with the live-death marker Zombie Violet (1:1000) (BioLegend, #423113). Cells were then pelleted to remove Zombie Violet and resuspended in 500 µl PBS.

Gating and analyses were done on the LSR II Fortessa at the Cytometry Facility (University of Zürich) using the 561 nm laser with a 576/26 nm filter to detect mOrange2 expression. At least 10^6 cells were recorded for the analysis per condition.

---

**Supplementary Figure 1:**
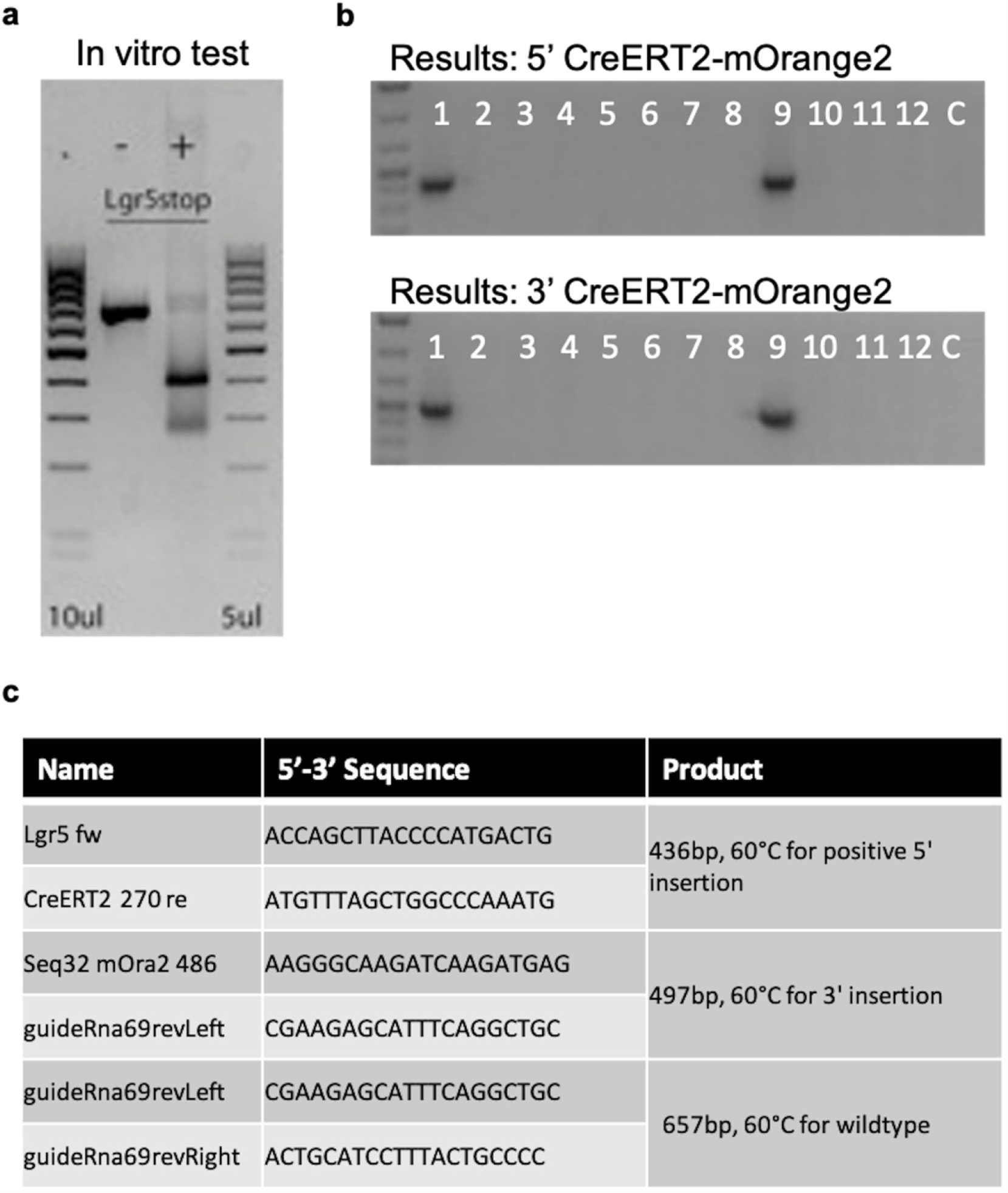
CRISPR/Cas9-mediated modification of murine Lgr5 locus. Lgr5 gRNA in vitro test to determine gRNA effectiveness on the target DNA. Left side - Cas9 protein, right side + Cas9 protein. b) Genotyping results for correct 5’ (top) and 3’ (bottom) integration of the CreERT2-mOrange2 cassette. c) Genotyping primer list with size product and PCR information

**Supplementary Figure 2:**
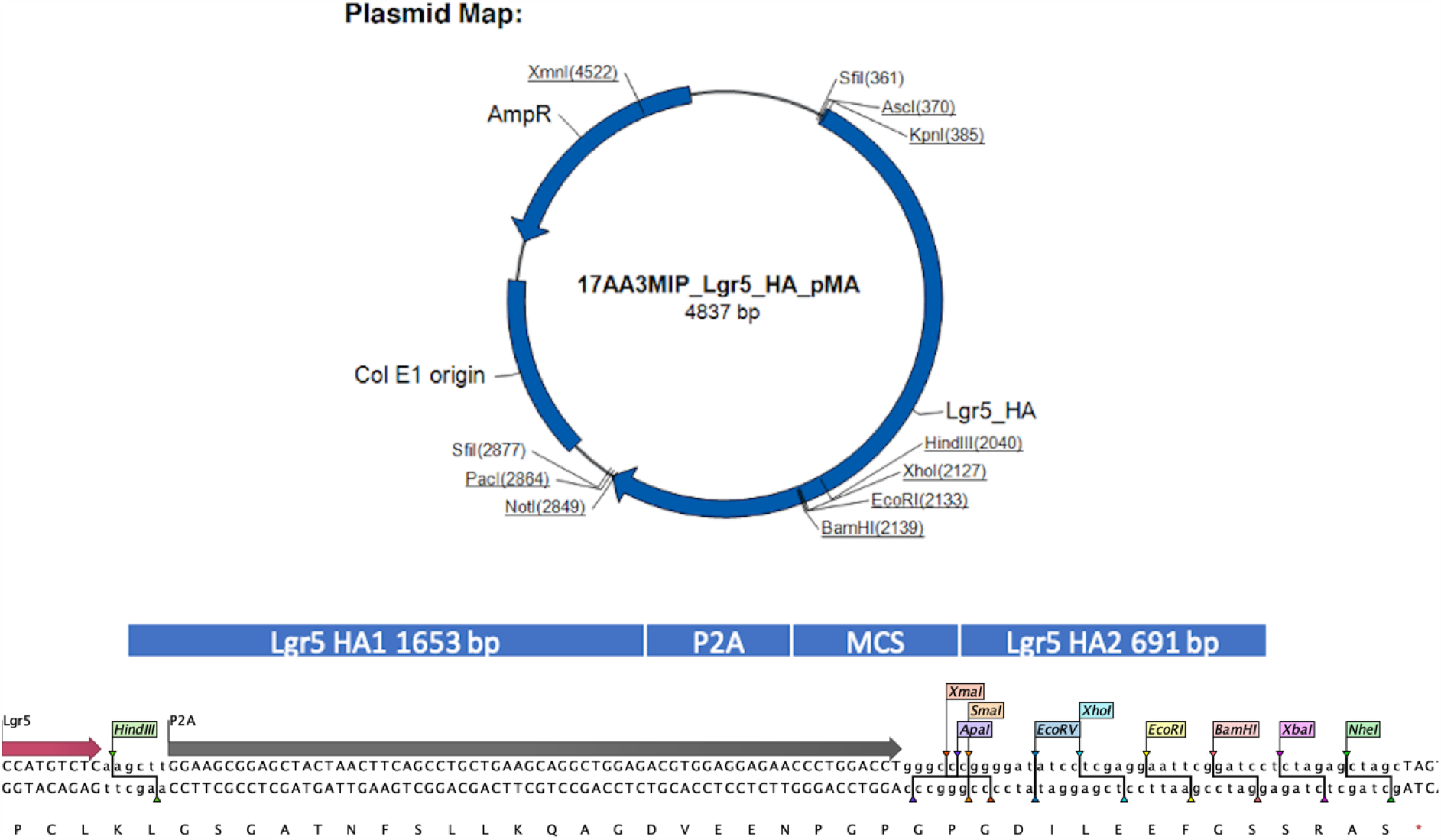
Schematic overview Lgr5 homology template. Plasmid overview is depicted top. The structure of the HRT is illustrated in the middle. The HRT harbours two homology arms (HA1 and HA2, sizes indicated in bp), which are flanking a P2A sequence and a multiple cloning site (MCS). The MCS was added to be able to clone different constructs in a fast and easy way. At the bottom the 3’ translational end of Lgr5 with the following P2A sequence and MCS are depicted in detail and the available restriction sites are indicated. The *CreERT2-P2A-mOrange2* cassette was cloned via XmaI and XbaI.

**Supplementary Figure 3:**
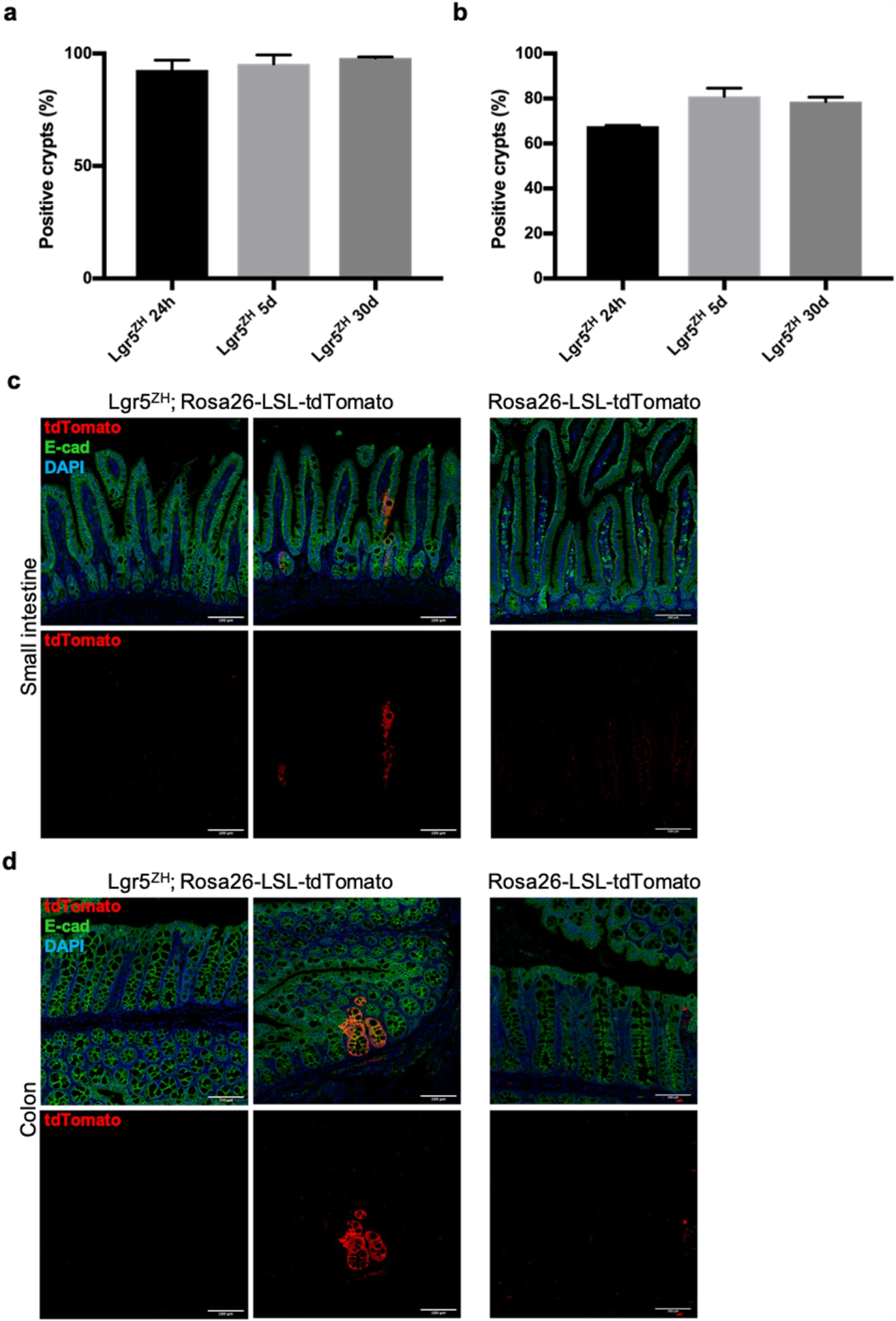
Quantification recombination efficacy within the crypts. Quantification of tdTomato^+^ crypts for the Lgr5^ZH^ allele for 24h, 5d and 30d in the small intestine a) and in the colon b). Cryps were considered positive when at least one cell was positive for tdTomato within the crypt. Minimum 5 independent stretches of crypts from proximal, mid and distal intestinal epithelium were analyzed per animal (n ≥ 2). c,d) Exemplary sections of uninjected Lgr5^ZH^; Rosa26-LSL-tdTomato animals in the small intestine, as well as in the colon. On rare occasions leaking of the CreERT2 recombinase could be observed. This was never found in Rosa26-LSL-tdTomato animals without the Lgr5^ZH^ driver.

